# Effects of Xiantao sewage treatment on microbial community structure of shallow groundwater in wetland

**DOI:** 10.1101/316802

**Authors:** Shu-Fen Song, Xiu-Fang Gao, Fan Yang

## Abstract

In order to solve the problem of urban domestic sewage treated by semi natural wetland method with high connectivity between surface water and underground water in the flood diversion channel, Whether there is a blank study on the pollution of shallow groundwater. The community structure and abundances of COD_Cr_, total nitrogen, total phosphorus and microorganism were added to the surface water. Environmental factors such as COD_Cr_, TN, TP, NH_4_^+^-N and microbial community and abundance indices were monitored through surface water and groundwater. In this study, physical and chemical indicators, microbial diversity and community structure of 12 water samples from Xiantao artificial wetland were studied by using the Illumina Miseq sequencing technique and the resulting microbial data were statistically analyzed in combination with environmental variables. The results showed that COD_Cr_ concentration had a very significant positive correlation with total microbial communities (r=0.530, P<0.01), NH_4_^+^-N and TN were significantly positively correlated (r=0.337 and 0.325, P<0.05). In addition, COD_Cr_ concentration was highly positively correlated with abundant groups (r=0.520, P<0.05), NH_4_^+^-N and TN were significantly positively correlated (r=0.325 and 0.304, P<0.05). For rare taxa, they might be more sensitive to the environment than their abundant groups. The relative abundance of the rare group is 0 at the depth of 10m, so we should carefully evaluate microbial reaction (rare group) environmental conditions in the future.

## Introduce

In recent years, the crisis in China’s groundwater has been highly concerned by the international community. In July 15, 2010, the News edition of the Nature^1^ magazine covered the whole page of the groundwater crisis in China (Qiu, 2010). In some areas, the storage of groundwater is decreasing at an alarming rate, and 90% of the water in the country has been polluted in varying degrees, of which 60% are seriously polluted.

According to the survey, in addition to landfill and gas station, urban sewage, industrial wastewater and agricultural wastewater from some large farms are also polluting in China’s urban groundwater (Xu, 2012). Some industrial and mining enterprises and large farms, driven by economic benefits, try to drain the sewage into the underground and think that the polluted water is “safe”. In fact, shallow groundwater and deep groundwater are only a relative concept, the distribution of groundwater is not uniform, and the absolute aquifers are not. The shallow groundwater and deep water are interconnected in many places, so the shallow groundwater will naturally flow into the deep layer.

Artificial wetland is an ecological wastewater treatment technology a new type of environmental protection (Yin, 2007), It has been successfully applied to storm flood, domestic sewage and eutrophication water purification (Kobayashi, *et al*., 2009; Iasur-Kruh, *et al*., 2010; Peralta, *et al*., 2012; Sánchez-Carrillo, *et al*., 2014).

The first example is to use the unused spillway semi natural wetland for urban sewage treatment, and the Xiantao wetland wastewater treatment plant system is made using the high water level surface water and the high connectivity water system of the groundwater. There are few studies on the influence of sewage treatment on groundwater by artificial wetland. The community and abundance of microbes in shallow groundwater under this condition, especially the effect of the cumulative results of pollutants on the groundwater under the long-term treatment conditions is almost not. The comprehensive study on the utilization of unused spillways, the treatment of domestic sewage by semi natural wetland, wetland method, the community and abundance of microbes in groundwater, the contamination of groundwater, the community and abundance of microbes in groundwater and the environmental factors of groundwater are the blank. The microbial diversity of surface water is one of the most important branches of water environment (Fu, *et al*., 2017; Guo, 2010). The physical and chemical properties of surface water vary with the conditions of space and light. However, the distribution of microorganisms in shallow groundwater under different depths is also a very important research topic. In this study, the samples not only have a wide range of samples, but also select three depths of 0.2m, 5M and 10m in the vertical direction. The community structure and abundance index of COD_Cr_, total nitrogen, total phosphorus and microbes are added to the surface water, and the environmental factors of surface water and groundwater (such as COD_Cr_, TN, TP, NH_4_^+^-N) and high throughput sequencing are monitored. Methods the microbial diversity was analyzed, and the structure and composition of microbial community in the shallow groundwater of domestic sewage, the relationship between the microbial community and the environmental factors were carried out by the wetland method. It can reflect the influence of the microbial community in the shallow groundwater during the operation of the wetland, and the correlation between the microbial community and the abundance change and the pollution level of the organic matter in the groundwater can provide the follow-up workers with the corresponding theoretical and practical support.

## 1. Materials and Methods

### 1.1 Introduction of Xiantao wetland method for domestic sewage treatment test

The Xiantao wetland experimental base is introduced in detail and the sampling azimuth reference Song (Song *et al*., 2015).

### 1.2 Determination of sampling and environmental factors of water samples

Methods of sampling and environmental factor determination refer to Song (Song *et al*., 2015).

### 1.3 Sequencing analyses

The processing method of raw data is obtained by referring to Song (Song *et al*., 2015).

## 2. Results

### 2.1 Analysis of physical and chemical indexes of wastewater

According to the state regulations, the treated sewage should reach the standard of pollutant discharge of GB 18918-2002 «municipal wastewater treatment plant», M1(400m) and M4(600m) were high concentration(COD_Cr_ > 120mg/L); M7(3000m) had achieved the two level standard of «municipal wastewater treatment plant»(100 mg/L < COD_Cr_ < 120 mg/L); and M2, M3, M5, M6, M8 and M9 were reach the first level standard (COD_Cr_ < 50 g/L). The TP of these samples were 0.02 –5.14mg/L, very significant positive correlation with COD_Cr_ (r=0.984, *P*<0.0001). Samples NH_4_^+^-N contents were 0.49–8.81mg/L, very significant positive correlation with COD_Cr_ (r = 0.987, *P*<0.0001). Samples TN contents were 1.13–19.17mg/L, very significant positive correlation with COD_Cr_ (r =0.983, *P*<0.0001) (**Table 1**). As can be seen from table 1, along with the level of wetland before treatment, the environmental factors gradually decreased. There is no phosphorus in the groundwater quality standard (GB/T 14848-2017). It is generally believed that the concentration of phosphorus in the groundwater is very low, but the phosphorus in the groundwater is present, and the amount of the phosphorus is large, and the pollution of phosphorus can not be ignored. The characteristics of the highly connected water system of surface water and groundwater are fully reflected here. From the surface water to the vertical direction of groundwater, that is, from surface water, groundwater depth of 5 meters, and buried depth of 10 meters, the environmental factors index is also gradually decreasing.

**Table 1.**
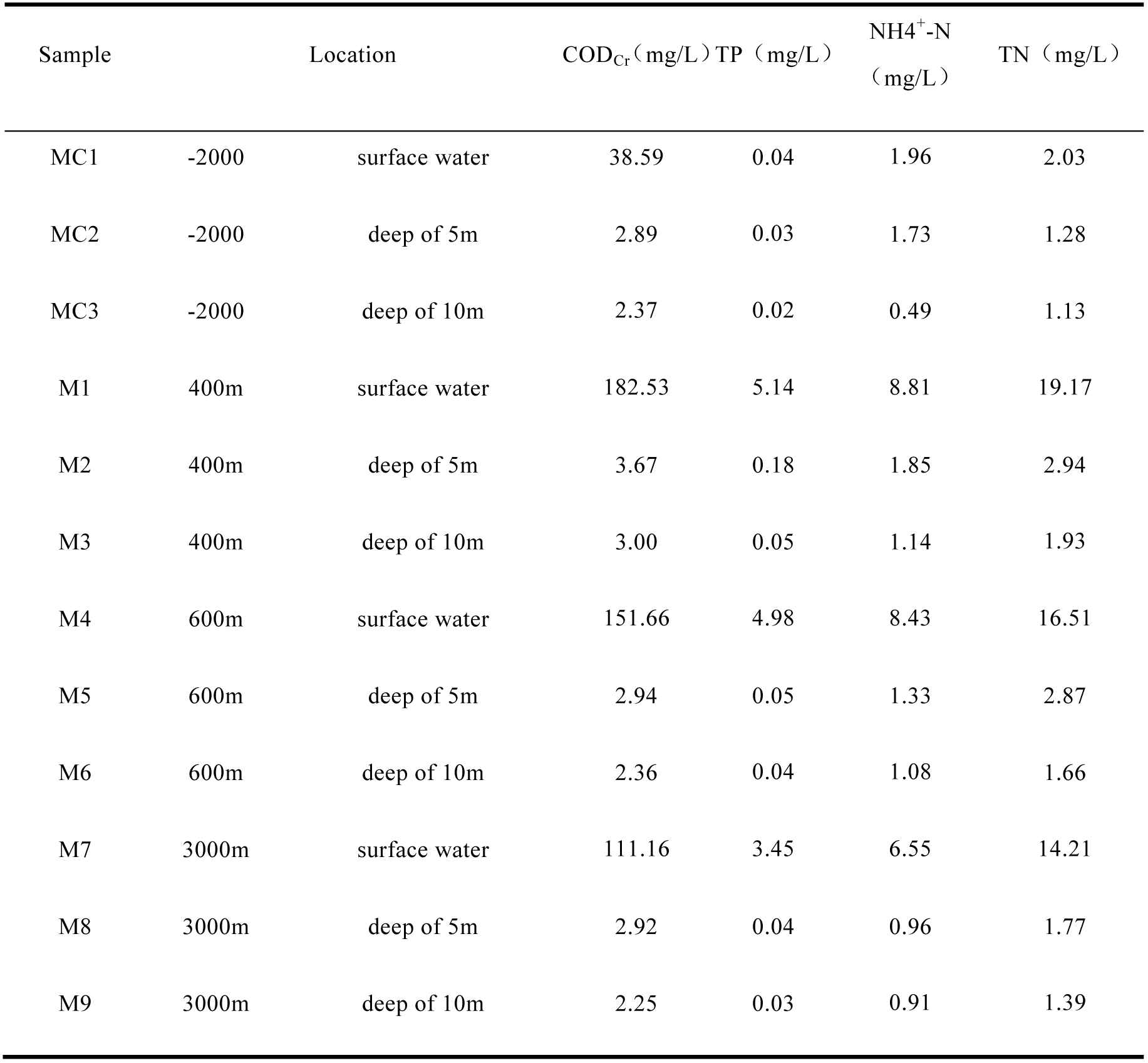
Detection results of surface water and groundwater in Constructed Wetland.

But from the horizontal direction of the wetland in the depth of 5 meters and 10 meters, the reduction is different.

### 2.2. Microbial diversity of the researched constructed wetland

A total of 80524 high-quality sequences with 5755-24785 sequences (mean=13454) and 726.9-1077.1 OTUs (mean=855) for twelve water samples were obtained. the diversity index, including Shannon (4.9-6.0), phylogenetic distance of a whole tree(48.1-68.5), and Chao 1 (2243.8–3291.9). All the calculated diversity indices in this study decreased with geographical position variation of azimuth of the studied samples (**Table 2**). The dominant phyla (the relative abundance of more than 1%) in the studied samples were *Proteobacteria*, *Bacteroidetes*, *Actinobacteria*, *Cyanobacteria*, *Verrucomicrobia*, *Firmicutes*, *Acidobacteria*, *Chloroflex* and *Euryarchaeota* (**Figure 1**). *Proteobacteria* is the most abundant phylum (more than 84% of total sequence reads).

**Figure 1.**
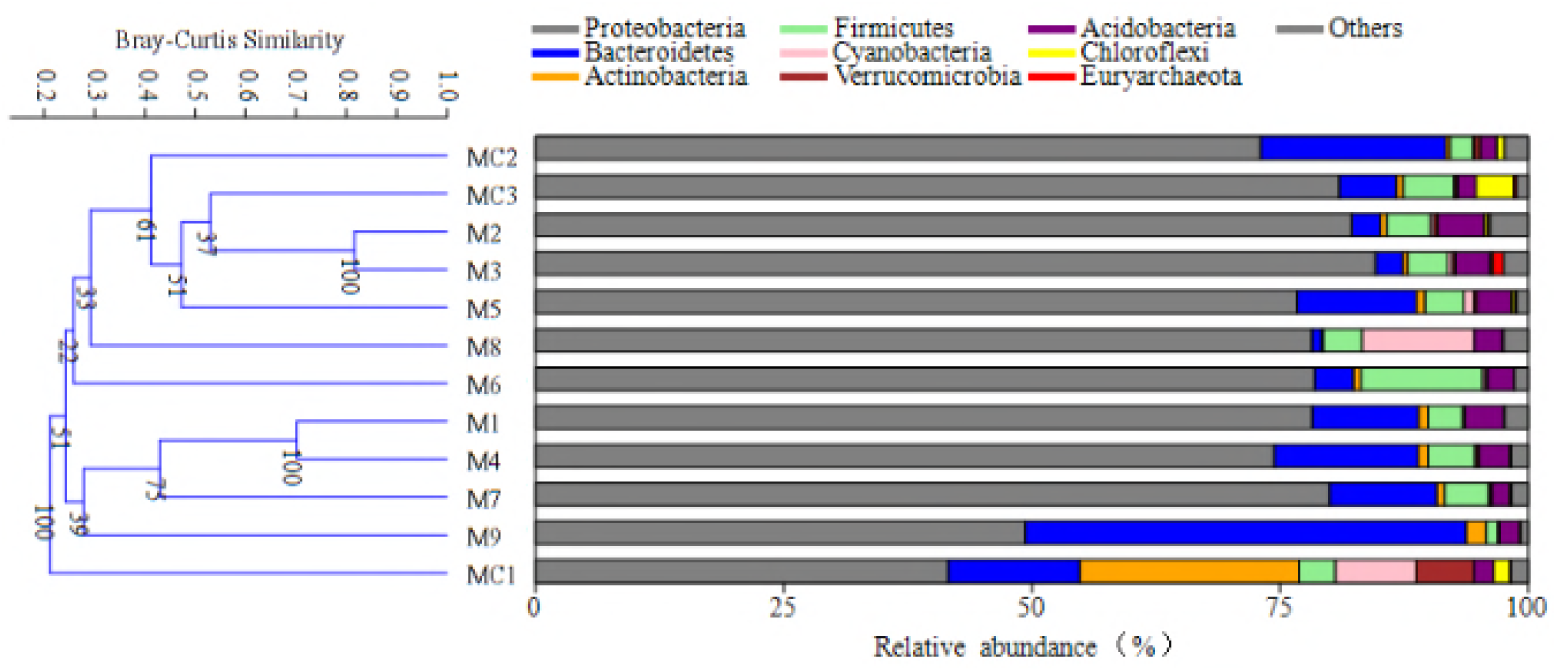
Bray-Curtis similarity-based cluster analysis (left) of structure of community in the studied samples and schematic figures (right) showing the frequencies of OTUs affiliated with major phyla in this study.

**Table 2.** Alpha-diversity of the surface water and groundwater in the studied Xiantao constructed wetland.

| sample | Total reads | Observed OTUs | Simpson | Shannon_ Wiener | PD_whole _tree | Good's Coverage | Chao1 |
| --- | --- | --- | --- | --- | --- | --- | --- |
| **MC1** | 18976 | 802.6 | 0.994 | 8.7 | 53.5 | 0.7 | 2393.2 |
| **MC2** | 21167 | 735.1 | 0.972 | 7.8 | 50.9 | 0.7 | 2306.8 |
| **MC3** | 16403 | 763.7 | 0.99 | 8.2 | 57.0 | 0.7 | 2336.8 |
| **M1** | 8237 | 966.7 | 0.987 | 8.7 | 59.3 | 0.6 | 3055.5 |
| **M2** | 11825 | 1077.1 | 0.992 | 9.2 | 68.5 | 0.6 | 3165.8 |
| **M3** | 7857 | 1046.8 | 0.991 | 9.0 | 66.6 | 0.6 | 3291.9 |
| **M4** | 11839 | 937 | 0.989 | 8.7 | 60.8 | 0.6 | 2638.8 |
| **M5** | 20513 | 897 | 0.993 | 8.8 | 55.3 | 0.6 | 3029.1 |
| **M6** | 7066 | 848.2 | 0.98 | 8.2 | 55.2 | 0.6 | 2898.5 |
| **M7** | 24785 | 730.8 | 0.975 | 7.6 | 57.6 | 0.7 | 2535.6 |
| **M8** | 7030 | 730.4 | 0.977 | 7.7 | 51.7 | 0.7 | 2419.5 |
| **M9** | 5755 | 726.9 | 0.985 | 8.0 | 48.1 | 0.7 | 2243.8 |

Among the retrieved OTUs, a total of 8-20 OTUs were classified as abundant OTUs. These abundant OTUs accounted for 5.63-15.15% of total OTUs and represented 64.66-80.85% relative abundance of sequence reads in the studied samples. In contrast, a total of 0-139 rare species were identified and they accounted for 0-64.19% of total OTUs and 0-6.98% relative abundance of sequence reads in the studied samples (**Table 3**). Most abundant OTUs belonged to *Proteobacteria* and *Bacteroidetes*, which the abundant OTUs accounted for 29.61-72.66% and 0-38.5% of total sequence reads in the studied samples (**Table 4**).

**Table 3.** Abundance estimates of the abundant and rare OTUs in the studied surface water and groundwater samples in this study.

| Sample | Abundant OTUs | Abundant OTU<br>relative abundance (%) | Rare OTUs | Rare OTU<br>relative abundance (%) |
| --- | --- | --- | --- | --- |
|  | (Percentage of<br>abundant OTUs/ total OTUs<br>in each sample) |  | (Percentage of<br>rare OTUs/ total OTUs<br>in each sample) |  |
| **MC1** | 20 (10.05%) | 67.96 | 92 (46.23%) | 5.27 |
| **MC2** | 17 (7.91%) | 77.93 | 138 (64.19%) | 6.98 |
| **MC3** | 16 (9.58%) | 79.76 | 96 (57.49%) | 5.75 |
| **M1** | 15 (10.42%) | 64.79 | 0 (0.00%) | 0 |
| **M2** | 8 (5.63%) | 66.82 | 36 (25.35%) | 3.27 |
| **M3** | 9 (7.69%) | 71.97 | 0 (0%) | 0 |
| **M4** | 17 (9.29%) | 64.66 | 37 (20.22%) | 2.47 |
| **M5** | 12 (6.45%) | 73.03 | 93 (50.00%) | 5.03 |
| **M6** | 13 (11.11%) | 74.34 | 0 (0.00%) | 0 |
| **M7** | 14 (6.01%) | 74.05 | 139 (59.66%) | 6.49 |
| **M8** | 12 (12.37%) | 80.85 | 0 (0.00%) | 0 |
| **M9** | 15 (15.15%) | 78.77 | 0 (0.00%) | 0 |

**Table 4.** Relative abundance of abundant OTUs within different phyla across the studied samples in this study.

| **Phylum** | Proteobacteria | Bacteroidetes | Actinobacteria | Cyanobacteria | Firmicutes | Verrucomicrobia | Chloroflexi | Euryarchaeota |
| --- | --- | --- | --- | --- | --- | --- | --- | --- |
| **MC1** | 29.61 | 5.95 | 19.71 | 5.95 | 1.06 | 4.61 | 1.06 | 0.00 |
| **MC2** | 63.30 | 14.64 | 0.00 | 0.00 | 0.00 | 0.00 | 0.00 | 0.00 |
| **MC3** | 72.66 | 2.15 | 0.00 | 0.00 | 1.29 | 0.00 | 3.66 | 0.00 |
| **M1** | 58.21 | 5.05 | 0.00 | 0.00 | 1.53 | 0.00 | 0.00 | 0.00 |
| **M2** | 65.37 | 0.00 | 0.00 | 0.00 | 1.45 | 0.00 | 0.00 | 0.00 |
| **M3** | 69.71 | 0.00 | 0.00 | 0.00 | 1.16 | 0.00 | 0.00 | 1.10 |
| **M4** | 54.80 | 8.39 | 0.00 | 0.00 | 1.47 | 0.00 | 0.00 | 0.00 |
| **M5** | 63.21 | 8.76 | 0.00 | 0.00 | 1.06 | 0.00 | 0.00 | 0.00 |
| **M6** | 66.31 | 0.00 | 0.00 | 0.00 | 8.03 | 0.00 | 0.00 | 0.00 |
| **M7** | 66.40 | 6.14 | 0.00 | 0.00 | 1.52 | 0.00 | 0.00 | 0.00 |
| **M8** | 68.14 | 0.00 | 0.00 | 11.18 | 1.53 | 0.00 | 0.00 | 0.00 |
| **M9** | 40.24 | 38.53 | 0.00 | 0.00 | 0.00 | 0.00 | 0.00 | 0.00 |

The studied shows that the water samples with similar physical and chemical tests have similar patterns of microbial structure. For example, cluster analysis revealed that the 12 samples are aggregated into two large clusters, of which the wetland surface water samples (M1, M4 and M7) are clustered into one cluster. The 5m deep and 10m deep water samples are divided into one cluster (except M9) (**Figure 2**). The horizonta direction Surface water of groundwater in Constructed Wetlands (MC1, M1, M4 and M7) were dominated by sequences affiliated with *Betaproteobacteria, Gammaproteobacteria*, *Epsilonproteobacteria*, *Actinobacteria*. The depth is 5 meters (MC2, M2, M5, and M8) were dominated by sequences affiliated with *Betaproteobacteria*, *Gammaproteobacteria*, *Bacteroidia*, *Melainabacteria*, *Sphingobacteriia*. The depth is 10 meters (MC3, M3, M6 and M9) were dominated by *Betaproteobacteria*, *Bacilli*, *Gammaproteobacteria*, *Alphaproteobacteria*, *Flavobacteriia*, *Sphingobacteriia*. the vertical of 400m (M1, M2 and M3) were dominated by sequences affiliated with *Betaproteobacteria.* 600m (M4, M5 and M6) were dominated by sequences affiliated with *Betaproteobacteria*, *Bacilli*, *Gammaproteobacteria*, *Alphaproteobacteria*, *Sphingobacteriia*. 3000m (M7, M8 and M9) were dominated by sequences affiliated with *Betaproteobacteria*, *Gammaproteobacteria*, *Flavobacteriia*, *Epsilonproteobacteria*, *Melainabacteria*, *Sphingobacteriia*. The contrast samples(MC1, MC2 and MC3) were dominated by *Betaproteobacteria*, *Gammaproteobacteria*, *Bacteroidia*, *Actinobacteria* (**Table 5**).

**Figure 2.**
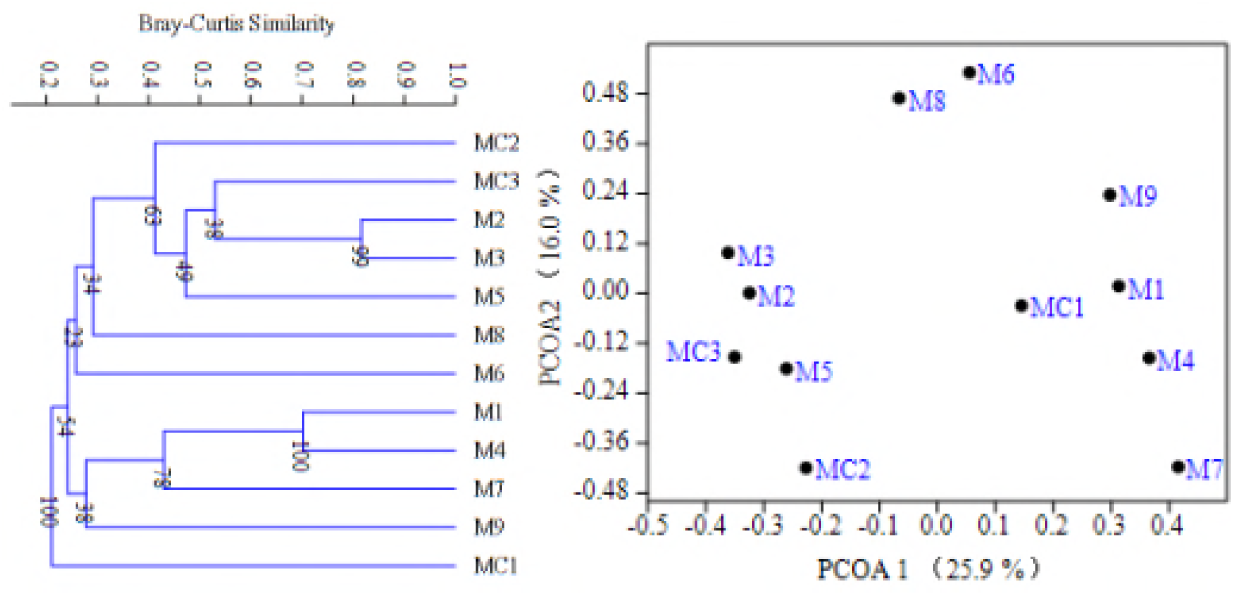
Clustering and principal coordinates analysis of total MCC among the studied samples based on Bray-Curtis similarity.

**Table 5.**
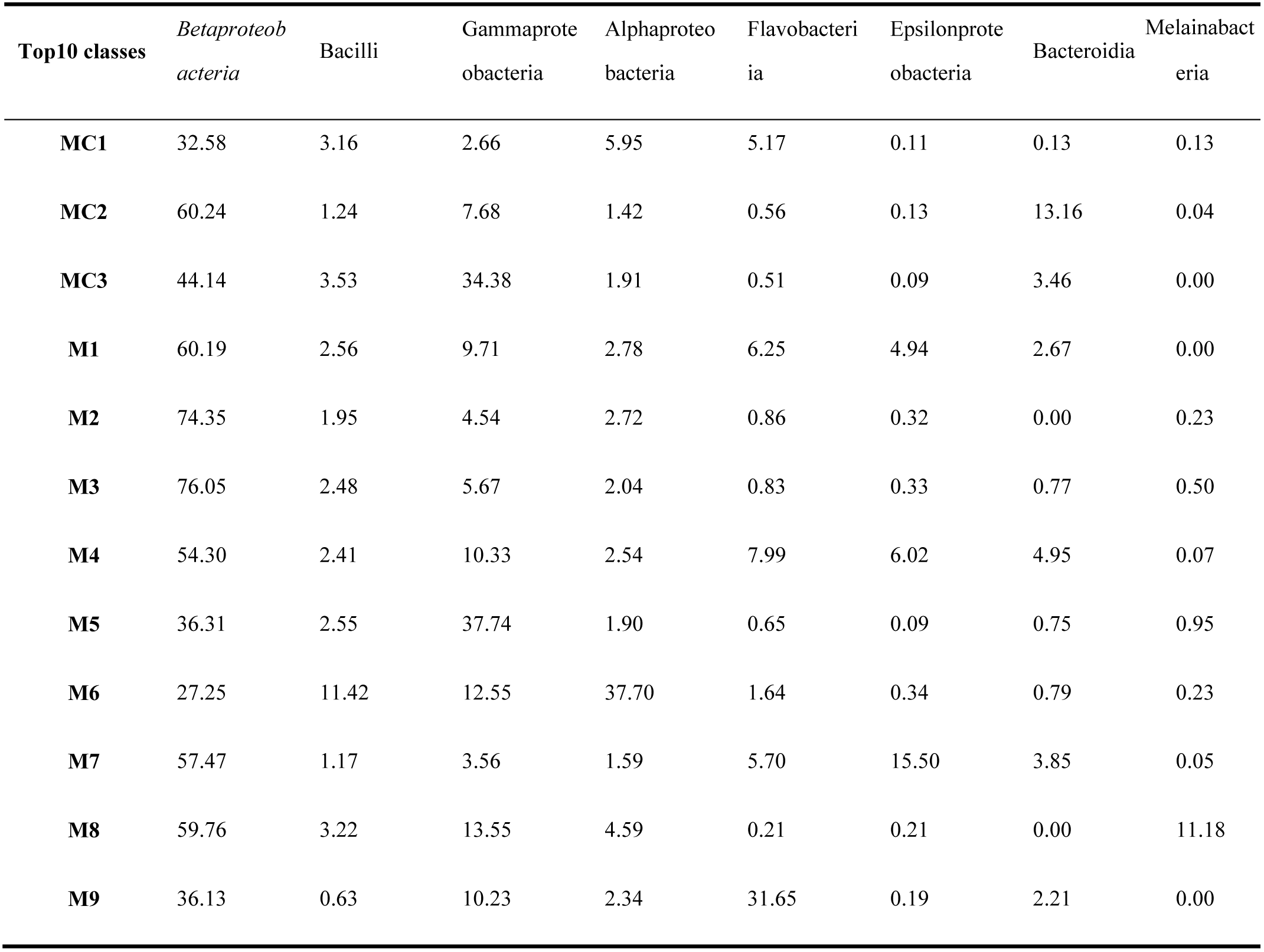
Relative abundance (%) of top 10 classes in the surface water and groundwater of the studied Xiantao constructed wetland.

| **Top10 classes** | *Betaproteobacteria* | Bacilli | Gammaproteobacteria | Alphaproteobacteria | Flavobacteriia | Epsilonproteobacteria | Bacteroidia | Melainabacteria |
| --- | --- | --- | --- | --- | --- | --- | --- | --- |
| **MC1** | 32.58 | 3.16 | 2.66 | 5.95 | 5.17 | 0.11 | 0.13 | 0.13 |
| **MC2** | 60.24 | 1.24 | 7.68 | 1.42 | 0.56 | 0.13 | 13.16 | 0.04 |
| **MC3** | 44.14 | 3.53 | 34.38 | 1.91 | 0.51 | 0.09 | 3.46 | 0.00 |
| **M1** | 60.19 | 2.56 | 9.71 | 2.78 | 6.25 | 4.94 | 2.67 | 0.00 |
| **M2** | 74.35 | 1.95 | 4.54 | 2.72 | 0.86 | 0.32 | 0.00 | 0.23 |
| **M3** | 76.05 | 2.48 | 5.67 | 2.04 | 0.83 | 0.33 | 0.77 | 0.50 |
| **M4** | 54.30 | 2.41 | 10.33 | 2.54 | 7.99 | 6.02 | 4.95 | 0.07 |
| **M5** | 36.31 | 2.55 | 37.74 | 1.90 | 0.65 | 0.09 | 0.75 | 0.95 |
| **M6** | 27.25 | 11.42 | 12.55 | 37.70 | 1.64 | 0.34 | 0.79 | 0.23 |
| **M7** | 57.47 | 1.17 | 3.56 | 1.59 | 5.70 | 15.50 | 3.85 | 0.05 |
| **M8** | 59.76 | 3.22 | 13.55 | 4.59 | 0.21 | 0.21 | 0.00 | 11.18 |
| **M9** | 36.13 | 0.63 | 10.23 | 2.34 | 31.65 | 0.19 | 2.21 | 0.00 |

### 2.3 Statistical analyses

Statistical analyses further corroborated the influence of different positions on the structure of community in the studied samples. Mantel test showed that structure of community of the treatment of surface water from domestic sewage by wetland method was correlated (*P*<0.01) with COD_cr_(r = 0.520) (**Table 6**). Furthermore, Bray-Curtis similarity of abundant microbial communities were significantly correlated (r=0.530) with treatment of surface water from domestic sewage by wetland method (**Figure 3**). Likewise, Mantel test indicated that the abundant structure of community were great significantly correlated to COD_cr_(r=0.530)(**Table 6**). In addition, Mantel tests also showed that three other indicators the total and abundant structure of community were significantly correlated with NH_4_^+^-N(r=0.325 and r=0.337), TN(r=0.304 and r= 0.325) (**Table 6**).

**Figure 3.**
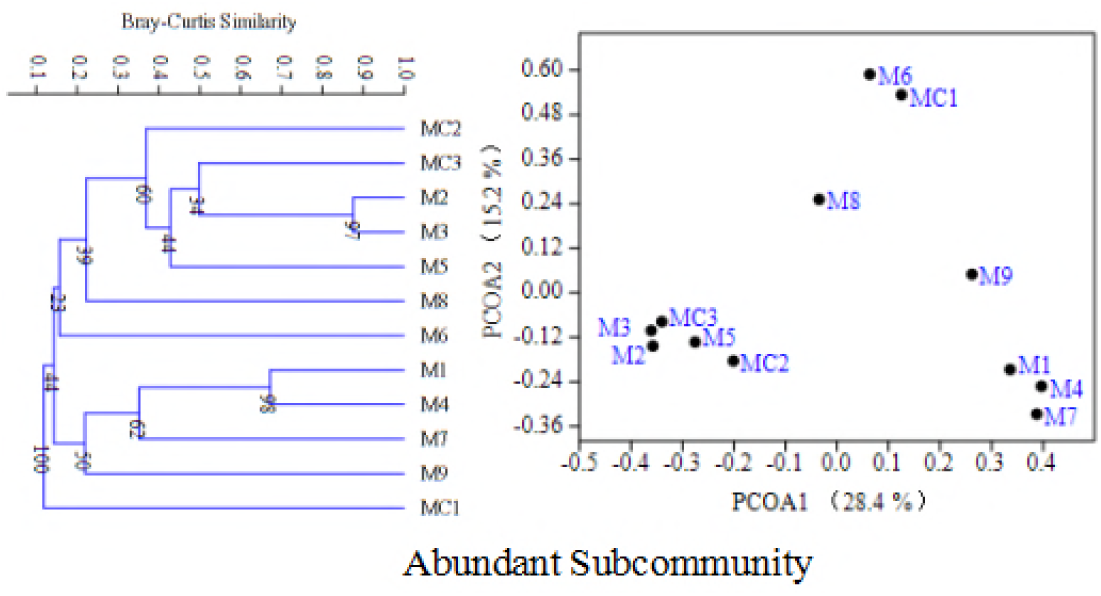
Cluster analyses and principal coordinates analyses of abundant structure of community among the studied samples based on Bray-Curtis similarity.

**Table 6.** Mantel test showing the correlation between structure of community similarity and environment parameters of the studied surface water and groundwater in this study.

|  | Abundant OTUs | All OTUs |
| --- | --- | --- |
| COD | 0.520** | 0.530** |
| TP | 0.206 | 0.232 |
| NH<sub>4</sub><sup>+</sup>-N | 0.325* | 0.337* |
| TN | 0.304* | 0.325* |

The Pearson's coefficients were calculated and their significances were tested based on 999 permutations.
\* $P < 0.05$ and \*\* $P < 0.01$ .

## 3. Discussion

The interaction of surface and groundwater covering almost in the process of water circulation, rainfall (Vidal, *et al*., 2008), interception (Schellekens, 2000), evaporation (Zhang, *et al*., 2008), infiltration (Yimer, *et al*., 2008), mathematical simulation and physical processes, and filtration (Fukada, *et al*., 2008), pollutants migration (Willhelm, *et al*., 1996) and other chemical processes, biological processes and the main contents of the study appear in the review article or some aspects of bioremediation for pollution.

In constructed wetlands, the removal of pollutants is mainly done through the physical, chemical and biological synergies of the matrix, microbes and plants in the wetland (Babatunde, *et al*., 2008). It is generally believed that biological action is the core factor of sewage purification in artificial wetland (Liang, *et al*., 2003). Some scholars through soil column simulation test, the effect of underground infiltration system on pollutant removal in sewage has been studied (Lance, *et al*., 1980). The results also confirm that the main removal effect in underground percolation system is biological action.

Combined with artificial wetland, the changes of microbial community in shallow groundwater in artificial wetland in the process of habitat restoration were compared and analyzed. It is found that geographical location is an important factor affecting the microbial diversity and community structure of the lake surface sediments. This finding is consistent with recent studies. Some scholars found that the number of microbes in fine sand decreased with the increase of sand depth (Liu, 2014). Some scholars found that the number of bacteria and fungi on the surface of the system was significantly higher than that of the lower layer. As the vertical height of the system deepened, the number of fungi decreased gradually (Zhou, *et al*., 2009). The total trend of bacteria and nitrite in the vertical direction is decreasing, and the number of microbes in the upper layer is more than that in the lower layer. The overall trend of bacteria and nitrite in the direction of water flow is also decreasing, that is, the front part is more than the middle and posterior parts (Fu, *et al*., 2005). Some scholars pointed out that in the same vertical subsurface flow wetland system, planting different plants also had an impact on the microbial community in the wetland, but they only had a significant impact on the 0~10 cm surface of the system, and the deeper microbial communities were basically similar (Sleytr, *et al*., 2009). In addition, in this study, experiments showed that the total microbial community structure was also significantly related to the NH_4_^+^-N and TN values of wetland, but lower than the correlation coefficient of COD_cr_ (r = 0.337, 0.325 VS R = 0.530, Table 6).

It is noteworthy that M9 and wetland surface water samples (M1, M4, M7), M6 and M8, M2, M3 and M5 three groups of samples gathered separately (**Figure 2**), indicating that some similar microorganisms may be shared between polluted area and water purification area. The reason for this phenomenon is that the concentration of pollutants in the surface water samples is too high to show the eutrophication state. Therefore, the number of microbes is large, and the concentration of pollutants decreases with the depth and direction of flow, and the nutrients are less, so the number of microbes is less.

It is also notable that the relative abundance of rare and abundant species of high surface water in the surface water treated in this study is less than that in the target area (**Table 3 and 4**), The rare communities showed more obvious reaction than the rich communities (which were proved by larger relative abundance), which indicated that rare taxa might have more restrictive distribution than the rich groups. Abundant groups can make use of abundant resources, thus having a very low probability of extinction and high probability of propagation. In addition, rare taxa may have less suitable niches in the 10m buried area, which are more vulnerable to environmental conditions than abundant taxa. Therefore, rare taxa may be responsive more sensitively to other environmental conditions than other rich taxa.

From the above research results, it can be seen that the geographical location is the most important factor affecting the microbial diversity and structure, whether it is a rich or rare community in the fairy peach artificial wetland. Rare groups are more sensitive to geographical location (possibly including other environmental conditions) than their abundant species.

## Acknowledgements

This research was supported by grants from the National Natural Science Foundation of China (Grant No. 41371464), Supported by the study abroad fund of Yangtze University.

We are very grateful to Professor He Ji Zheng of the Chinese Academy of Sciences, the center for ecological environment research.

